# Computational and experimental investigation of biofilm disruption dynamics induced by high velocity gas jet impingement

**DOI:** 10.1101/824532

**Authors:** Lledó Prades, Stefania Fabbri, Antonio D. Dorado, Xavier Gamisans, Paul Stoodley, Cristian Picioreanu

## Abstract

Experimental data showed that high-speed micro-sprays can effectively disrupt biofilms on their support substratum, producing a variety of dynamic reactions such as elongation, displacement, ripples formation and fluidization. However, the mechanics underlying the impact of high-speed turbulent flows on biofilm structure is complex in such extreme conditions, since direct measurements of viscosity at these high shear rates are not possible using dynamic testing instruments. Here we used computational fluid dynamics simulations to assess the complex fluid interactions of ripple patterning produced by high-speed turbulent air jets impacting perpendicular to the surface of *Streptococcus mutans* biofilms, a dental pathogen causing caries, captured by high speed imaging. The numerical model involved a two-phase flow of air over a non-Newtonian biofilm, whose viscosity as a function of shear rate was estimated using the Herschel-Bulkley model. The simulation suggested that inertial, shear and interfacial tension forces governed biofilm disruption by the air jet. Additionally, the high shear rates generated by the jet impacts coupled with shear-thinning biofilm property resulted in rapid liquefaction (within milliseconds) of the biofilm, followed by surface instability and travelling waves from the impact site. Our findings suggest that rapid shear-thinning in the biofilm reproduces dynamics under very high shear flows that elasticity can be neglected under these conditions, behaving the biofilm as a Newtonian fluid. A parametric sensitivity study confirmed that both applied force intensity (i.e. high jet-nozzle air velocity) and biofilm properties (i.e. low viscosity, low air-biofilm surface tension and thickness) intensify biofilm disruption, by generating large interfacial instabilities.

**IMPORTANCE:** Knowledge of mechanisms promoting disruption though mechanical forces is essential in optimizing biofilm control strategies which rely on fluid shear. Our results provide insight into how biofilm disruption dynamics is governed by applied forces and fluid properties, revealing a mechanism for ripples formation and fluid-biofilm mixing. These findings have important implications for the rational design of new biofilms cleaning strategies with fluid jets, such as determining optimal parameters (e.g. jet velocity and position) to remove the biofilm from a certain zone (e.g. in dental hygiene or debridement of surgical site infections), or using antimicrobial agents which could increase the interfacial area available for exchange, as well as causing internal mixing within the biofilm matrix, thus disrupting the localized microenvironment which is associated with antimicrobial tolerance. The developed model also has potential application in predicting drag and pressure drop caused by biofilms on bioreactor, pipeline and ship hull surfaces.

## INTRODUCTION

During recent years close attention has been paid to the mechanical behavior of biofilms when subjected to high shear turbulent flows, because its direct application in developing methods for biofilm removal. Fluid-biofilm patterns such as formation of migratory ripples and surface instabilities, transition to fluid-like behavior and stretching to finally break off the biofilm have been described (1–5). Furthermore, the viscoelastic character of biofilms when exposed to turbulent flows has been reported (6–8), hypothesizing that generated instabilities enhance mass transfer into biofilms (9, 10).

Several biofilm “constitutive” models, mathematical models which describe the mechanical properties of material and how a material will respond to mechanical forces, have been developed considering viscoelastic properties and biofilm-fluid interaction, as reviewed by different authors (11–14). Biofilm deformation and detachment under laminar flow conditions have been modeled using the phase field approach (15, 16) and the immersed boundary method (17). While the movement of biofilms streamers (18), and the deformation of simple-shaped biofilm under high-speed flow (19) have been described using fluid-structure interaction models, implemented in commercial software packages (i.e. COMSOL Multiphysics and ANSYS). Small biofilm deformations have also been modelled considering the biofilm as a poroelastic material, compressing when exposed to laminar flows (20, 21). The effect of parallel air/water jets on the surface morphology of very viscous biofilms has been investigated numerically (3, 22), proposing the characterization of the biofilm rippling as Kelvin-Helmholtz instabilities. However, the interactions between biofilm and perpendicular impinging turbulent jets flow, and how the turbulent impacts disrupt biofilm properties have not yet been described numerically. Therefore, a numerical investigation of the biofilm rippling patterns generated by turbulent jets could help clarifying the mechanics behind the observed biofilm disruption. Computational fluid dynamics (CFD) is a useful tool for predicting the behavior of fluid flow and fluid-fluid interactions which might not be easily possible through experimentation. CFD models have been developed to describe air-jet impingements into water vessels (23–25), using the volume of fluid (VOF) method to track the interface position between fluids. VOF method has also been used to predict the wall shear stress produced by turbulent flows over biofilms (26), and to characterize the biofilm removal by impinging water droplets (27). Nevertheless, both the turbulence effect on the growth surface and the biofilm as a distinct dynamic phase have not been considered. Thus, a CFD model for the biofilm rippling under turbulent jets should include: (i) multiphase flow with biofilm and air/water both as moving phases; (ii) an accurate tracking of the biofilm-fluid interface with VOF; (iii) a reliable treatment of fluid dynamics at the biofilm growth surface (i.e. near-wall treatment) and at the biofilm-fluid interface (e.g. turbulent damping correction); and (iv) the biofilm phase as a non-Newtonian fluid, with liquefaction due to shear thinning at high shear rates.

The present study was aimed at developing a CFD model to characterize the observed dynamic rippling patterns of *Streptococcus mutans* biofilms exposed to high velocity air-jet perpendicular impingements. To this goal, two-dimensional (2D) axisymmetric CFD simulations were performed to describe the behavior of air-impinged biofilms, considering turbulence and near-wall treatment. The non-Newtonian biofilm was examined under high shear rates, to reveal the mechanisms driving its disruption under air-jet impacts. The model was used to study the influence of parameters, such as biofilm viscosity at rest and at the predicted high shear rates, biofilm thickness, nozzle-jet velocity and air-biofilm surface tension, on biofilm cohesiveness, deformation and disruption.

## MATERIALS AND METHODS

### Biofilm growth

*S. mutans* biofilms UA159 (ATCC 700610) were grown for 72 hours on glass microscope slides at 37 °C and 5% CO_2_ in a brain-heart infusion (Sigma-Aldrich, St. Louis, MO) supplemented with 2% (wt/vol) sucrose (Sigma-Aldrich, St. Louis, MO) and 1% (wt/vol) porcine gastric mucin (Type II, Sigma-Aldrich, St. Louis, MO). After the growth period, the biofilm-covered slides were gently rinsed in 1% (wt/vol) phosphate-buffered saline solution (Sigma-Aldrich, St. Louis, MO) and placed in Petri dishes (28). Biofilm thickness was determined by fixing untreated samples with 4% (wt/vol) paraformaldehyde and staining with Syto 63 (Thermo Fisher Scientific, UK). Subsequently, three random confocal images were taken on three independent replicate biofilm slides, measuring a thickness of 51.8±4.9 μm by COMSTAT software (27), as previously described (25), thus we took the biofilm thickness of *L_b_*=55 μm in the simulations.

### Biofilm perpendicular air jet impingement

An air jet generated from a piston compressor (ClassicAir 255, Metabo, Nürtingen, Germany) impinged on biofilm samples at a 90° angle. Experiments were performed in triplicate. The compressor tip (internal diameter of 2 mm) was held at a 5 mm distance from the biofilm. The air jet impingement was recorded at 2000 frames per second with a high-speed camera (MotionPro X3, IDT Vision, Pasadena, US), placed to record the back view of the biofilm-covered microscope slide. The average air jet velocity exiting the nozzle (v=41.7±1.5 m·s^-1^) was measured with a variable area flow meter. To estimate Reynolds number for the biofilm (Re_b_) flowing along the substratum, the biofilm thickness (*L_b_*) was used as the characteristic length and the biofilm density (ρ_b_) was assumed to equal to water. The variation of biofilm displacement velocity (*v_b_*) and biofilm viscosity (*η*) variables had greater effect in the Re_b_. At the highest shear stresses, the biofilm behaved as if it was completely liquefied to water with *η*=0.001 Pa·s (see *Results* section, Figure 3) and moved with a maximum velocity *v_b_*≈0.2 m·s^-1^. Under these conditions Re_b_=11 indicating laminar flow. Considering the viscosity and density of air at 20°C and 1 atm, and the characteristic length equal to the nozzle tip diameter, an estimated maximum Reynolds number of the air-jet (Re_a_) was 5600, which is in the turbulent regime (29).

### Data post-processing

Fast Fourier transform (FFT) was used to determine the dominant period (T) and dominant frequency (f) of the ripple patterns in the *S. mutans* biofilm formed during exposure to the air jet. Frames from the experimental high-speed video and data exported from the simulated results were post-processed with a MATLAB script for the FFT analysis.

For the experimental high-speed videos, the ripples wavelength (λ_R_), defined as the distance between two reverse peaks, was measured using NIH ImageJ as previously described (28). Briefly, videos were converted to stacks and λ_R_ was measured with the “plot profile” function. For the simulated data, λ_R_ was computed post-processing the biofilm surface contours using MATLAB. The ripples velocity (u_R_, the distance the wave travels in a given time) was calculated as u_R_ = f·λ_R_ = λ_R_/T.

### Numerical model

The general assumptions made in the development of the numerical model were:

1. The gas (air) and liquid (biofilm) phases are incompressible (Mach number below 0.3 for the gas phase).
2. A uniform velocity profile, constant in time, leaves the compressor nozzle.
3. The flow is symmetric with respect to the vertical axis (around nozzle middle).
4. The free gas jets are in the turbulent regime (due to calculated Re_a_).
5. The initial biofilm is a thin layer with constant thickness.
6. The biofilm is characterized by non-Newtonian fluid shear-thinning (Herschel-Bulkley model) and density equal to that of water.

#### Model geometry

The schematic representation of the experimental set-up and the computational domain used to analyze the air-jet impingements over biofilm thin layer are shown in Figure 1A and Figure 1B, respectively. A 2D axisymmetric computational domain represented the lateral view of the jet impact over biofilm, slicing the actual experimental set-up. The domain length and height were L_x_=15 mm and L_y_=5 mm respectively, with the biofilm initial thickness *L_b_*=0.055 mm.

**Figure 1.**
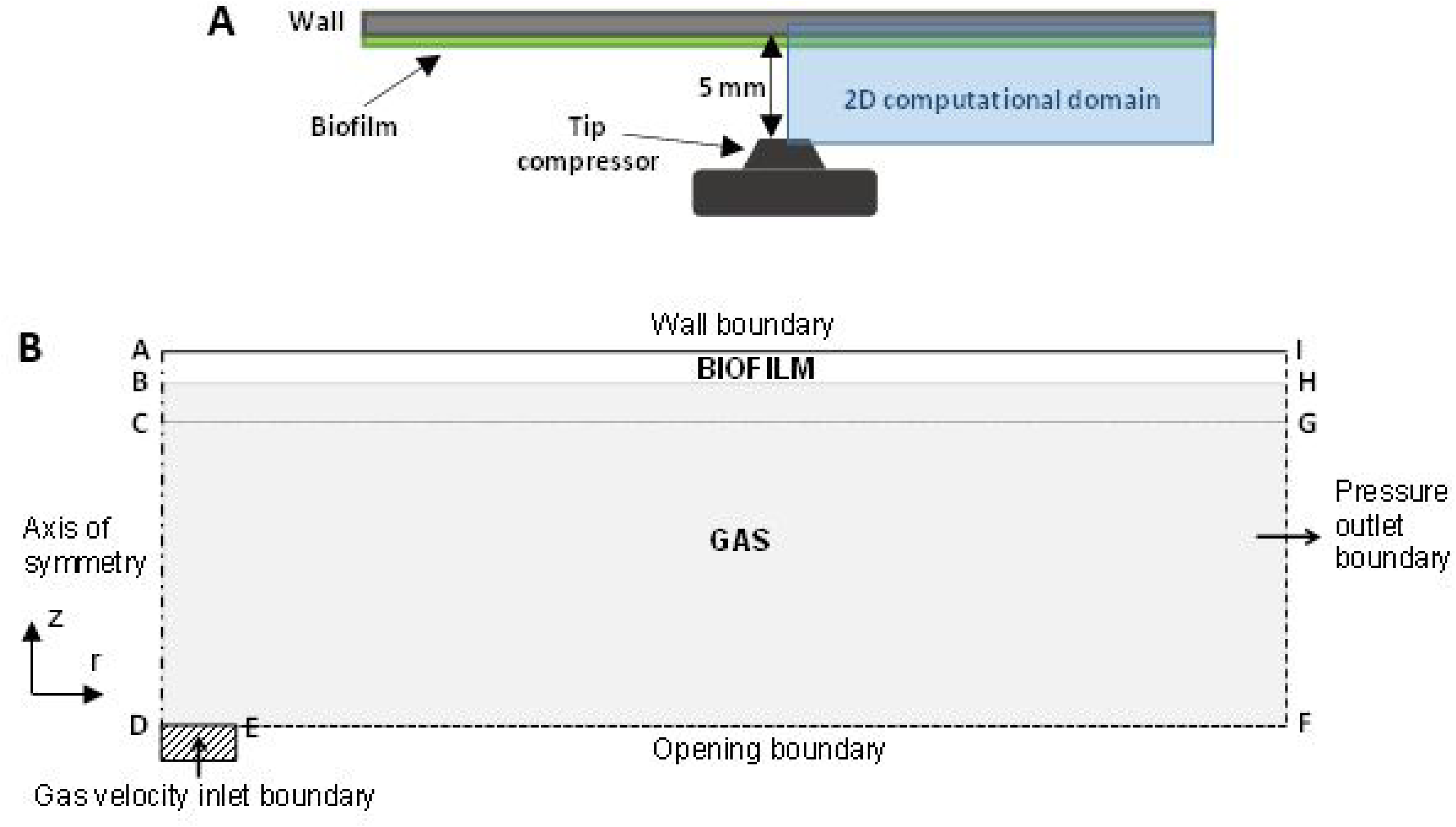
Experimental set-up (A) and two-dimensional axisymmetric model (B) with radial (r) and axial (z) directions and the boundary conditions. A no-slip and zero-turbulence *wall* was imposed on the boundary AI, representing the glass microscope slide, the substratum that the biofilm was grown on. The air *inlet* was established on DE and a *pressure outlet* condition was set on the boundary FI. A *symmetry axis* was used along on the boundary AD, and the boundary EF was *open* to the atmosphere.

##### Governing equations

###### Mass and momentum conservation

The momentum conservation eq.(1) is coupled with the continuity eq.(2)

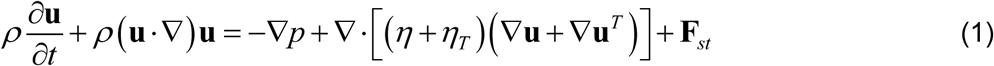

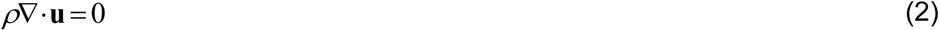

solved for the local velocity vector **u** and pressure *p*. **F***_st_* is the force arising from surface tension effects. The fluid density *ρ* and dynamic viscosity *η* were calculated by the VOF method in each control volume. *η_T_* is the turbulent viscosity, resulting from the k-ω turbulence model (see *Turbulence model* section). The interface between fluids (i.e. air and biofilm) was tracked with a robust coupling between level-set and VOF methods, as implemented in the ANSYS Fluent software (30).

###### Turbulence model

An examination of Reynolds-averaged Navier–Stokes (RANS) modeling techniques recommends the shear-stress transport (SST) k-ω model instead of the standard k-ε model, since it can describe better fluid flow in impinging jets within reasonable computational effort (29). The SST k-ω model incorporates a blending function to trigger the standard k-ω model in near-wall regions and the k-ε model in regions away from the wall. The turbulence kinetic energy, k, and the specific dissipation rate, ω, are obtained from the transport equations including the convection and viscous terms, together with terms for production and dissipation of k and ω and cross-diffusion of ω. A user-defined source term for ω, representing the turbulence damping correction, was added to correctly model the flows in the interfacial area. Turbulence damping was needed because otherwise the large difference in physical properties of biofilm and air phases would create a large velocity gradient at the interface, resulting in unrealistically high turbulence generation (31). See Supplementary Information for more details.

###### Biofilm viscosity

The Herschel-Bulkley model (32) was used to characterize the dynamic viscosity of *S. mutans* biofilms, representing previously observed shear-thinning non-Newtonian behavior. The dynamic viscosity *η* (Pa s) is inversely related to the shear rate γ (s^-1^) and proportional to the shear stress *σ* (Pa):

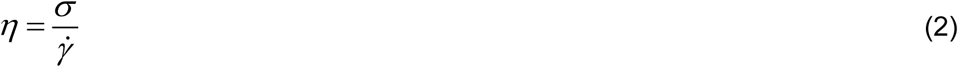

while the shear stress depends on the shear rate:

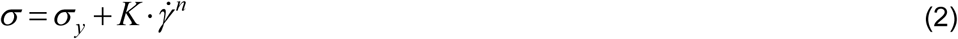

with *σ_y_* the yield stress (Pa), *K* the consistency index (Pa·s) and *n* the flow behavior index. Since the calculated shear rates from CFD were so high that experimental data could not be obtained (orders of magnitude higher than obtainable with ordinary rheometers) we extrapolated using data from dynamic viscosity sweeps to determine the complex viscosity of *S. mutans* biofilms (33) and the dynamic viscosity of heterotrophic biofilms determined in (34), by modifying Herschel-Bulkley model parameters and evaluating numerically several viscosity curves from (eq.(2)). See results for more details.

###### Boundary and initial conditions

In the computational domain (Figure 1B), a symmetry axis was used on the boundary AD, with radial velocity component and normal gradients equal to zero. The boundary EF was open to the atmosphere, depending on the mass balance. A zero gauge pressure outlet condition was set on FI. The air inlet was on DE (half-nozzle size), with a fully-developed velocity profile. The inlet turbulent energy, k, was computed as:

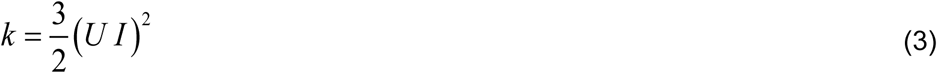

with *U* the mean flow velocity and *I* the turbulence intensity defined as *I* = 0.16 · *Re*^−1/8^ (23), and Reynolds number *Re* defined with velocity *U* and nozzle diameter. The specific turbulent dissipation rate, *ω*, was:

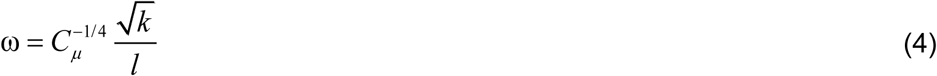

with the empirical constant *C_μ_* = 0.09 and *l* the turbulent length scale, assumed 7% of the nozzle diameter.

A no-slip and zero-turbulence wall was imposed on the biofilm substratum (boundary AI), with near-wall formulation to represent precisely the wall-bounded turbulent flow in the region, including the buffer layer and viscous sublayer. The *y+-insensitive near-wall treatment* (30) was used here, where based on the dimensionless wall distance of the first grid cell (y^+^), the linear and logarithmic law-of-the-wall formulations were blended. To resolve the viscous sublayer, the first grid cell needed to be at about y^+^≈1, also near the free surface (31). In the two-phase flow, the biofilm phase was initialized as a thin layer with constant thickness over the substratum, being several orders of magnitude more viscous than the air. The biofilm behaved initially like a solid, requiring resolving the viscous sublayer from the air-biofilm interface, instead of from the wall as usually done.

In the initialization step, the values for *k* and *ω* were computed using equations (3) and (4), from velocity *U* in the inlet and characteristic length *l* = 1 mm, and the volume fraction of the biofilm phase was set to 1 in the region ABHI (Figure 1B).

##### Model solution

###### Meshing

A uniform mesh of prism cells was defined in the domain, with maximum size h_x_ x h_y_ of 50 μm x 17 μm, with a refined mesh in the region ACGI (minimum size 15 μm x 0.4 μm) to satisfy the requirement y^+^≈1 near walls and in the free surface. A mesh growth rate no higher than 1.2 (30) was used between the refined sub-domain and the remaining computational domain, leading to ∼450000 mesh cells. Mesh details are shown in Supplementary Information **Figure S1**.

###### Solvers

The mathematical model was implemented into the commercial fluid dynamics software ANSYS Fluent (Academic Research, Release 17.2). The governing equations were discretized using a second-order upwind scheme in space and first-order implicit in time, with pressure staggering option interpolation (PRESTO) and pressure-implicit splitting of operators (PISO) for the pressure-velocity coupling. The free surface deformation was tracked with the geo-reconstructed scheme. Transient simulations ran with a maximum time step set to 10^-7^ s for stable transient solutions. A total time of 20 ms was simulated in each run to reach a quasi-stationary solution.

###### Simulation plan

Two sets of simulations were carried out with the two-phase model. The first set (runs 1-7) was performed for model calibration, where the biofilm viscous properties were evaluated according to the experimental data. Experimental parameters, such as the measured jet velocity (*v*) and biofilm thickness (*L_b_*), were used in this set with different non-Newtonian viscosities (*η*) (i.e. estimated viscosity curves, EVC) and two surface tensions (*γ*). The second set (runs 8-12) was performed for sensitivity analysis, evaluating the effects of the inlet jet velocity and the biofilm thickness on the biofilm rippling response. Table 1 shows an overview of the numerical simulations.

**Table 1.** Overview of numerical simulations of jet impingements at velocity *v* on biofilm layer with thickness *L_b_*, surface tension *γ* and viscosity *η*.

| Run | 1 | 2 | 3 | 4 | 5 | 6 | 7 | 8 | 9 | 10 | 11 | 12 |
| --- | --- | --- | --- | --- | --- | --- | --- | --- | --- | --- | --- | --- |
| $\eta$ (Pa·s) | EVC<sub>3</sub> | EVC<sub>3</sub> | EVC<sub>1</sub> | EVC<sub>1</sub> | EVC<sub>2</sub> | EVC<sub>2</sub> | EVC<sub>4</sub> | EVC<sub>3</sub> | EVC<sub>3</sub> | EVC<sub>3</sub> | EVC<sub>3</sub> | EVC<sub>3</sub> |
| $\gamma$ (mN·m<sup>-1</sup>) | 36 | 72 | 36 | 72 | 36 | 72 | 36 | 36 | 72 | 36 | 72 | 36 |
| $L_b$ (μm) | 55 | 55 | 55 | 55 | 55 | 55 | 55 | 55 | 55 | 55 | 55 | 27.5 |
| $v$ (m·s<sup>-1</sup>) | 41.7 | 41.7 | 41.7 | 41.7 | 41.7 | 41.7 | 41.7 | 20 | 20 | 60 | 60 | 41.7 |

## RESULTS

### Experimental results

Figure 2A depicts image frames recorded at different times (5, 10, 15 and 20 ms) with high-speed camera during perpendicular air jet impingement. The movies showed how the air-jets first generated a clearing in the biofilm at the impingement site followed by the formation of surface instabilities which rapidly spread radially. The disrupted area grew for approximately 200 ms until it stabilized with a diameter of approximately 1.5 cm. A movie showing the process is in the Supplementary Information **Video S1**. After ∼350 ms the ripples died out when the biofilm had flowed to the cleared space edge.

**Figure 2.**
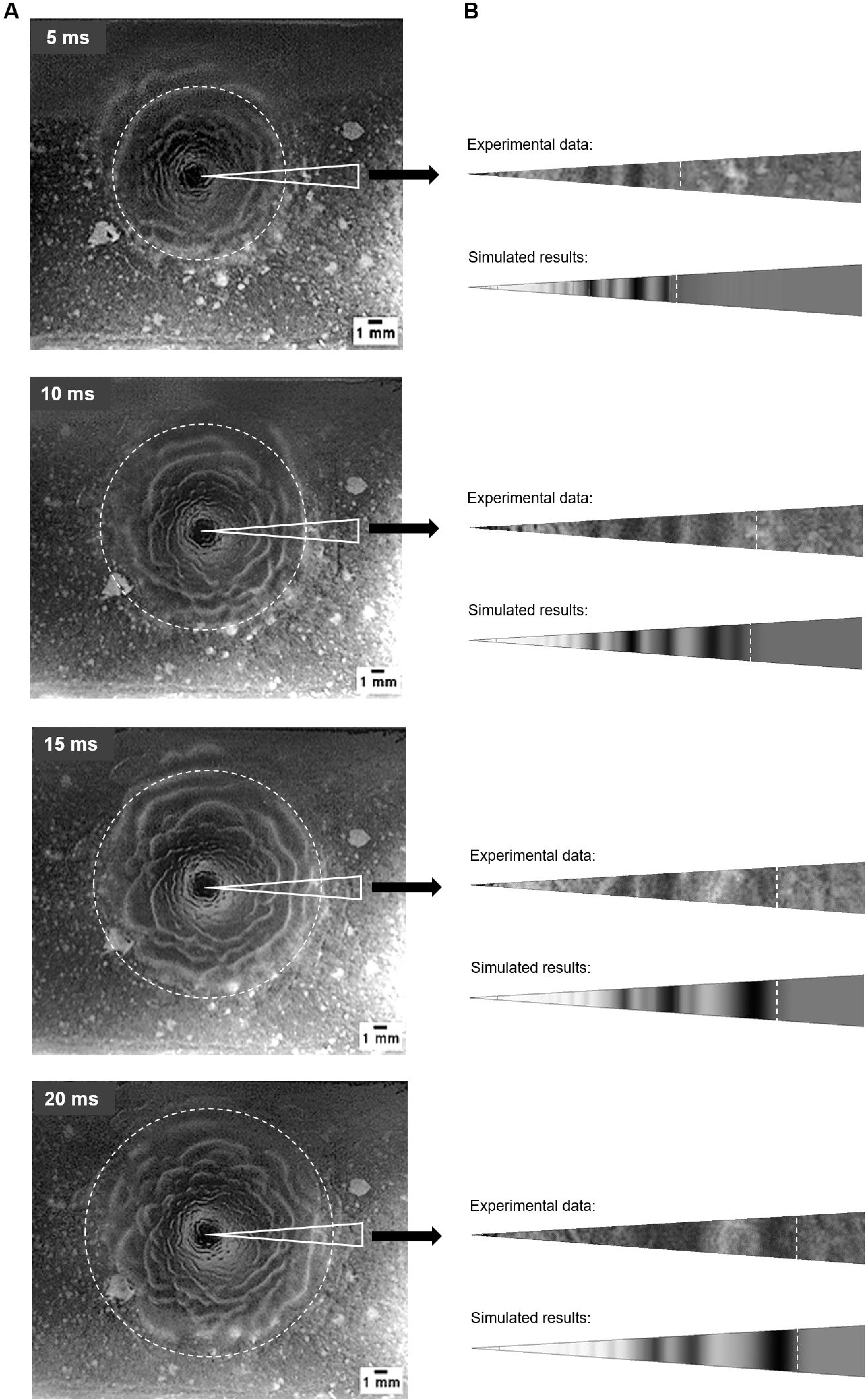
(A) Image frames at 5, 10, 15 and 20 ms from the high-speed movie. Thicker biofilm ripples and cell clusters outside the ripple zone are light and the background slide surface is dark. (B) Comparison of measured and simulated biofilm displacement (right column) within the marked sector, with EVC_3_ and *γ*=36 mN·m^-1^. The front position of the advancing ripples is indicated by the white dashed line. The gray scale in the simulations shows the local biofilm thickness, which is correlated with the wave amplitude. The high-speed recording of the jet impingement experiment is available in Supplementary Information **Video S1**. The animation of the simulated ripple formation is presented in the Supplementary Information **Video S2**.

### Model verification with experimental data

#### Biofilm viscosity assessments

The shear-thinning character of *S. mutans* biofilms has been previously demonstrated by measuring the complex viscosity (33), however no data is reported about their characterization as non-Newtonian fluids. The model development required determining the dynamic viscosity of *S. mutans*-based biofilms at much higher shear rates (10^4^–10^6^ s^-1^) than commonly reported for other biofilms (up to 2000 s^-1^). Shear rate measurements in this range are impracticable with normal rheometric systems which can achieve shear rates of around 10^4^ s^-1^. Microchannels containing MEMS pressure sensors have been used to achieve almost 10^5^ s^-1^ (35), however even this is still an order of magnitude less than the predicted shear rate experienced by the biofilm in our experiments. Thus we were forced to extrapolate the values used in the model from existing data. The complex viscosity can be related to the dynamic viscosity by the empirical Cox-Merz rule (36) (i.e. both viscosity measures should be identical at comparable observation time-scales). However, some discrepancies in the Cox-Merz rule have been reported in samples with gel characteristics such as polysaccharides and biofilms by obtaining larger values for the complex viscosities (34, 37, 38), probably due to physical and chemical interactions present in these samples (32).

Assuming that the dynamic viscosity should be lower than the complex viscosity, parametric sweeps were performed to evaluate the dynamic viscosity using the Herschel-Bulkley model from eqns.(2) and (2)). As an example, four estimated viscosity curves (EVC_1_, EVC_2_, EVC_3_, and EVC_4_) were represented together with the experimental reference values in Figure 3. EVC_1_ corresponded to the highest dynamic viscosity, while EVC_4_ was the lowest viscosity curve. To reproduce the observed liquefaction behavior in the movies (9), the Herschel-Bulkley curves were adjusted to bend asymptotically to that of water viscosity at very high shear rates with the reasoning that water represents the lowest possible limit for a completely broken down hydrogel. However, in reality the biofilm viscosity is expected to be higher than that of water since even if completely mixed will contain cells and EPS components. The Herschel-Bulkley parameters, and the shear rate thresholds at which biofilm viscosity reached the viscosity of water are listed in Table 2.

**Figure 3.**
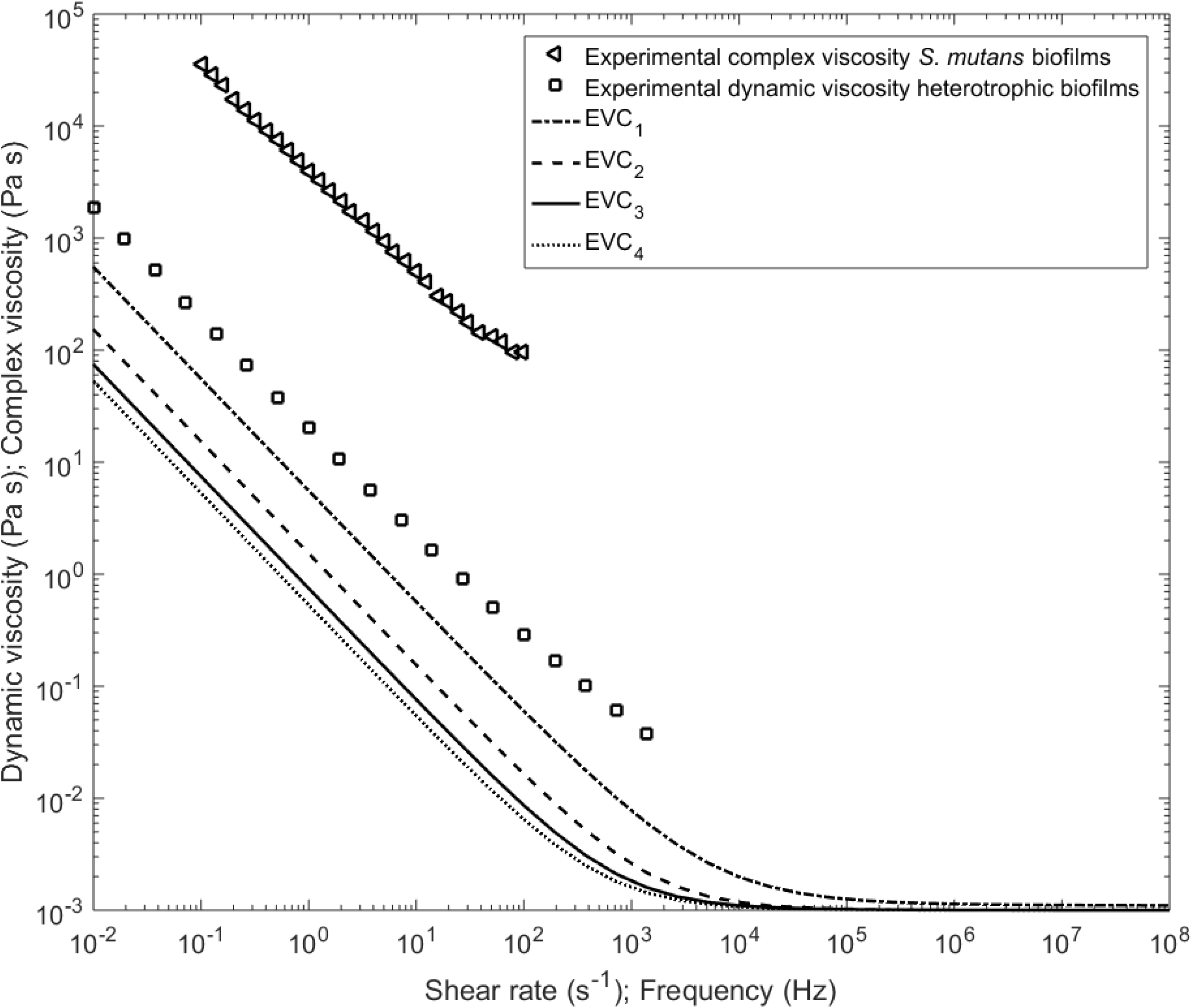
Dependency of biofilm viscosity on shear rate. Experimental dynamic viscosity for heterotrophic biofilms (squares) function of shear rate (34) and the complex viscosity of *S. mutans* (triangles) function of frequency (33). Solid lines represent estimated viscosity curves (EVC) with different parameter values as in Table 2.

**Table 2.**
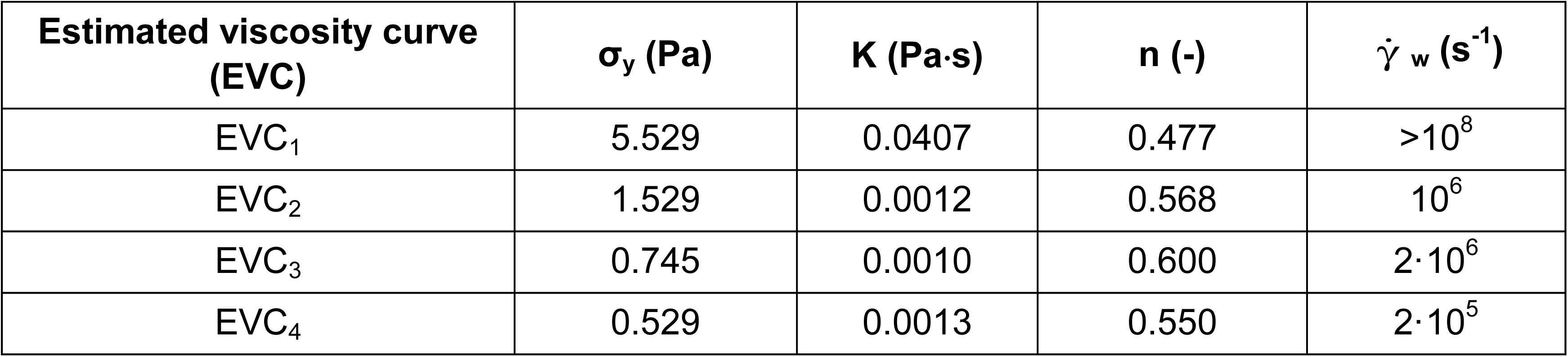
Rheological parameters of Herschel-Bulkley model (yield stress *σ_y_*, fluid consistency index *K* and flow behavior index *n*) and threshold shear rates at which biofilm viscosity reached water viscosity (_w_) for the four estimated viscosity curves.

| Estimated viscosity curve (EVC) | $\sigma_y$ (Pa) | $K$ (Pa·s) | $n$ (-) | $\dot{\gamma}_w$ (s<sup>-1</sup>) |
| --- | --- | --- | --- | --- |
| EVC<sub>1</sub> | 5.529 | 0.0407 | 0.477 | $>10^8$ |
| EVC<sub>2</sub> | 1.529 | 0.0012 | 0.568 | $10^6$ |
| EVC<sub>3</sub> | 0.745 | 0.0010 | 0.600 | $2 \cdot 10^6$ |
| EVC<sub>4</sub> | 0.529 | 0.0013 | 0.550 | $2 \cdot 10^5$ |

#### Computed results

For the experimental air inlet velocity *v*=41.7 m·s^-1^, the biofilm response was simulated with different viscosity curves (Table 2) and two surface tension values (*γ*) of air-biofilm interface (*γ*=72 mN·m^-1^ and *γ*=36 mN·m^-1^). The former is for air-water *γ* at 20 °C, considering the biofilm matrix as more than 90% water (39), and the latter, smaller value is from Koza et al. (40) since the amphiphilic nature of EPS can have a surface-active effect (41), altering the air-water *γ*. The image frames taken during the biofilm disruption at different times were compared with simulation results. The estimated viscosity curve EVC_3_ with *γ*=36 mN·m^-1^ best matched the experimental data as illustrated in Figure 2B, with respect to the distance reached by the travelling wave front and also the position of several ripple maxima and minima thicknesses (dark/light areas in the simulation results). An animation of the simulated ripple formation is presented in the Supplementary Information **Video S2**. Figure 4 depicts the biofilm surface contours over time for cases simulated with EVC_3_ and *γ*=72 mN·m^-1^ (left) or *γ*=36 mN·m^-1^ (right), from the early cavity formation (Figures 4 A,D) until the deformation wave damping (Figures 4 C,F). An animation of the simulated biofilm rippling can be found in Supplementary Information **Video S3**. A lower surface tension clearly intensified the disruption and the formation of surface instabilities, which were qualitatively analyzed at 4 and 20 ms (Figure 4). Ripples began to form from 3 and 5 mm at 4 and 20 ms, respectively. At early times (4 ms) the cavity width was ∼2 mm and depth reached >80% of the initial biofilm thickness. Later, at 20 ms the cavity width extended to ∼4 mm and the depth reached almost to the biofilm substratum, creating a zone cleared of biofilm with a radius of about 2 mm. The disruption caused a biofilm deceleration of ∼0.025 m·s^-2^.

**Figure 4.**
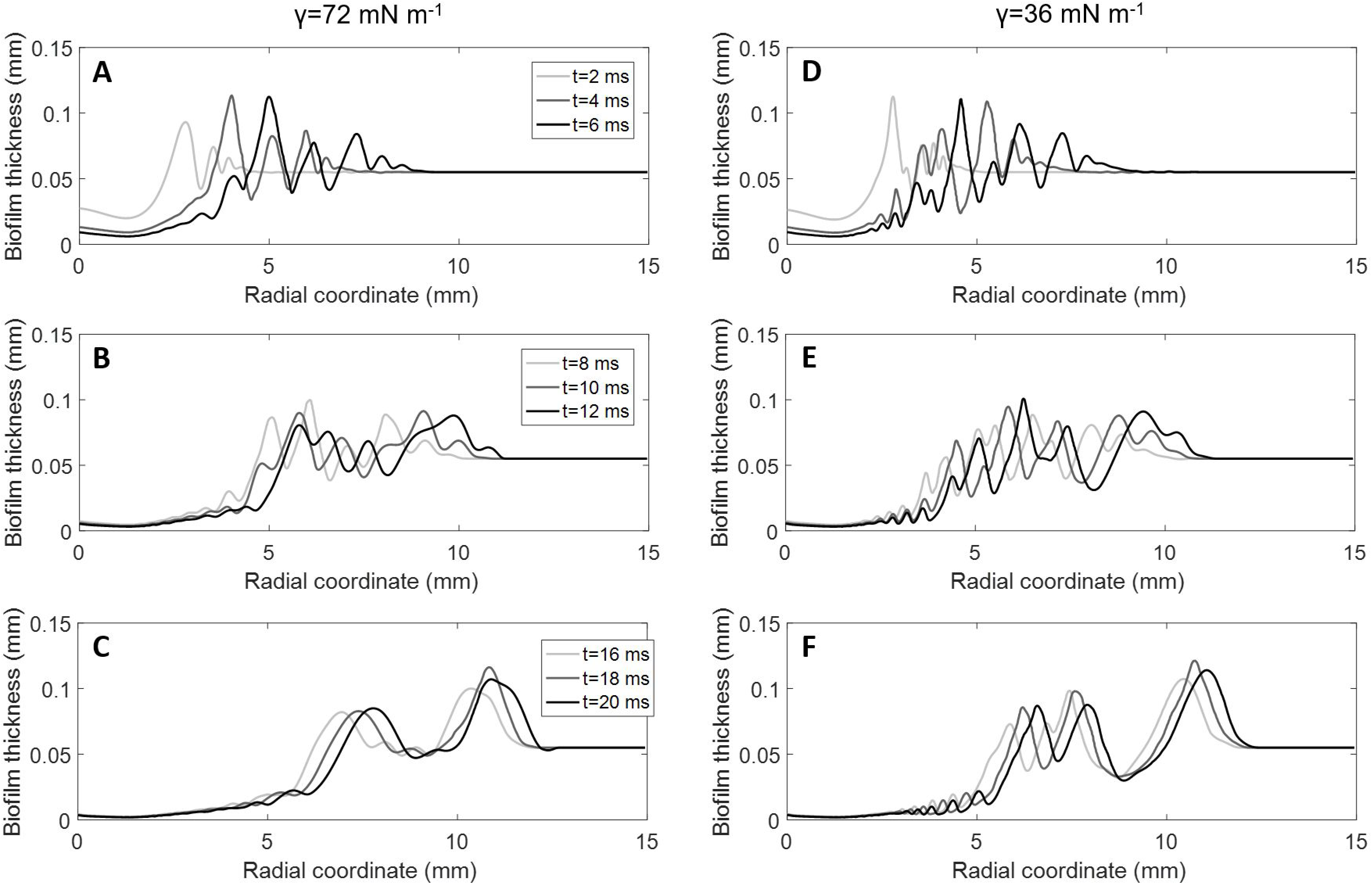
Simulated changes of biofilm thickness in time as a function of radial distance from the point of impingement (from 2 to 20 ms) for viscosity model EVC_3_ and two surface tensions: (A-C) *γ*=72 mN·m^-1^ and (D-F) *γ*=36 mN·m^-1^. An animation of the simulated biofilm rippling can be found in Supplementary Information **Video S3**.

By tracking the position of the advancing front of the ripples, the biofilm displacement was determined as defined as the maximum distance travelled by the advancing front of the ripples over the underlying biofilm support at a given time (5). From the movies an average displacement was computed from the front positions in eight radial directions at each time (Figure 2), whereas the biofilm displacement in the simulations was computed in one radial direction, because of the axial symmetry of the computational domain. Figure 5 compares the experimental and simulated displacements, where biofilm responses were computed with EVC_3_ and two surface tensions. Initially the disruption front moved quickly, but slowed down and reached a steady value after about 20 ms, trends reflected in both experiments and model. It seems that the lower surface tension allowed a faster displacement (i.e. less opposing force to the air stream) and better fit of the experimental data, but the differences were still too small to be considered significant.

**Figure 5.**
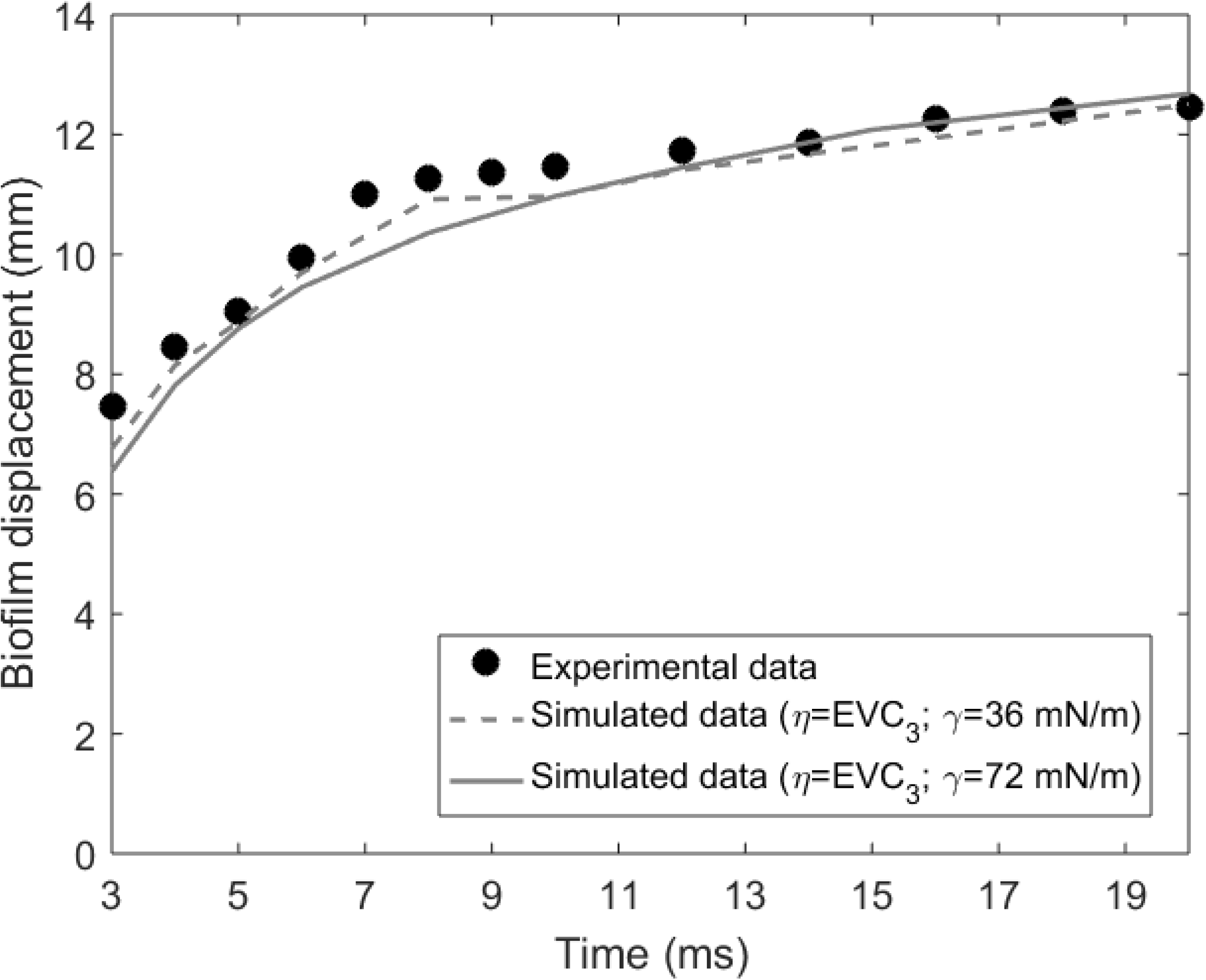
Experimental (symbols) and simulated data (lines) of biofilm displacement as a function of jet exposure time.

A simulated sequence of the air-jet impingement over the biofilm in the lateral view showing fluid-biofilm interaction is presented in Figure 6 (for an animation see Supplementary Information **Video S4**). Initially, the air flow flowed faster than the biofilm, forming ripples on the biofilm surface, until a steady-state was reached after 20 ms, generating a disrupted zone with ∼13 mm radius. The velocity field shows the high velocity around the air nozzle and continuously decreasing velocities as the air flows radially along the biofilm surface far from the impact zone. The air flow loses its kinetic energy as it expands radially, and at a certain distance from the center the shear would be too weak to deform the biofilm anymore, therefore a steady-state was reached.

**Figure 6.**
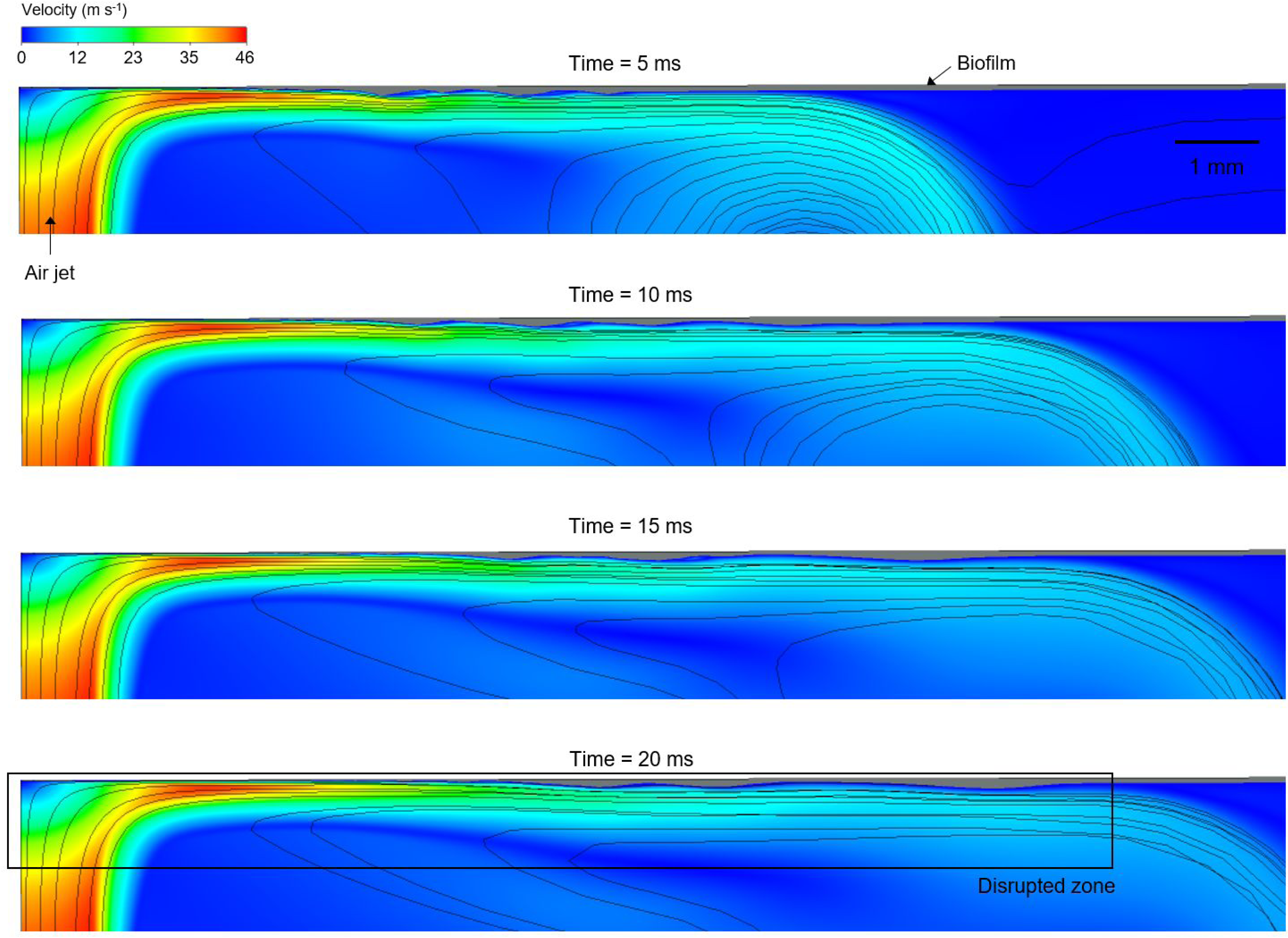
Simulated sequence (5, 10, 15 and 20 ms) of the air-jet impingement over the biofilm, for *η*=EVC_3_ and *γ*=36 mN·m^-1^. Velocity magnitude of the air flow is represented by the colored surface, while the biofilm is the gray area on the top side. Air streamlines and flow directions are also displayed. (For interpretation of the references to color in this figure legend, the reader is referred to the Web version of this article.) An animation of the simulated biofilm rippling can be found in Supplementary Information **Video S4**.

A sequence of the air shear rate and the biofilm dynamic viscosity distributions are presented in Figure 7 (for an animation see Supplementary Information **Video S5**), and the pressure and velocity components profiles are depicted in **Figure S3** and **Figure S4** respectively (see Supplementary Information). The simulations showed high shear rates (∼10^6^-10^7^ s^-1^), which directly produced high shear stresses (∼10^4^ Pa) and relative pressure (∼1400 Pa) in the air-jet impact zone during air-jet exposure. Significantly, the high shear rates generated on the air-biofilm interface rapidly reduced the biofilm dynamic viscosity from the initial value (7500 Pa·s) to that of water (0.001 Pa·s), suggesting complete liquefaction of the biofilm had occurred.

**Figure 7.**
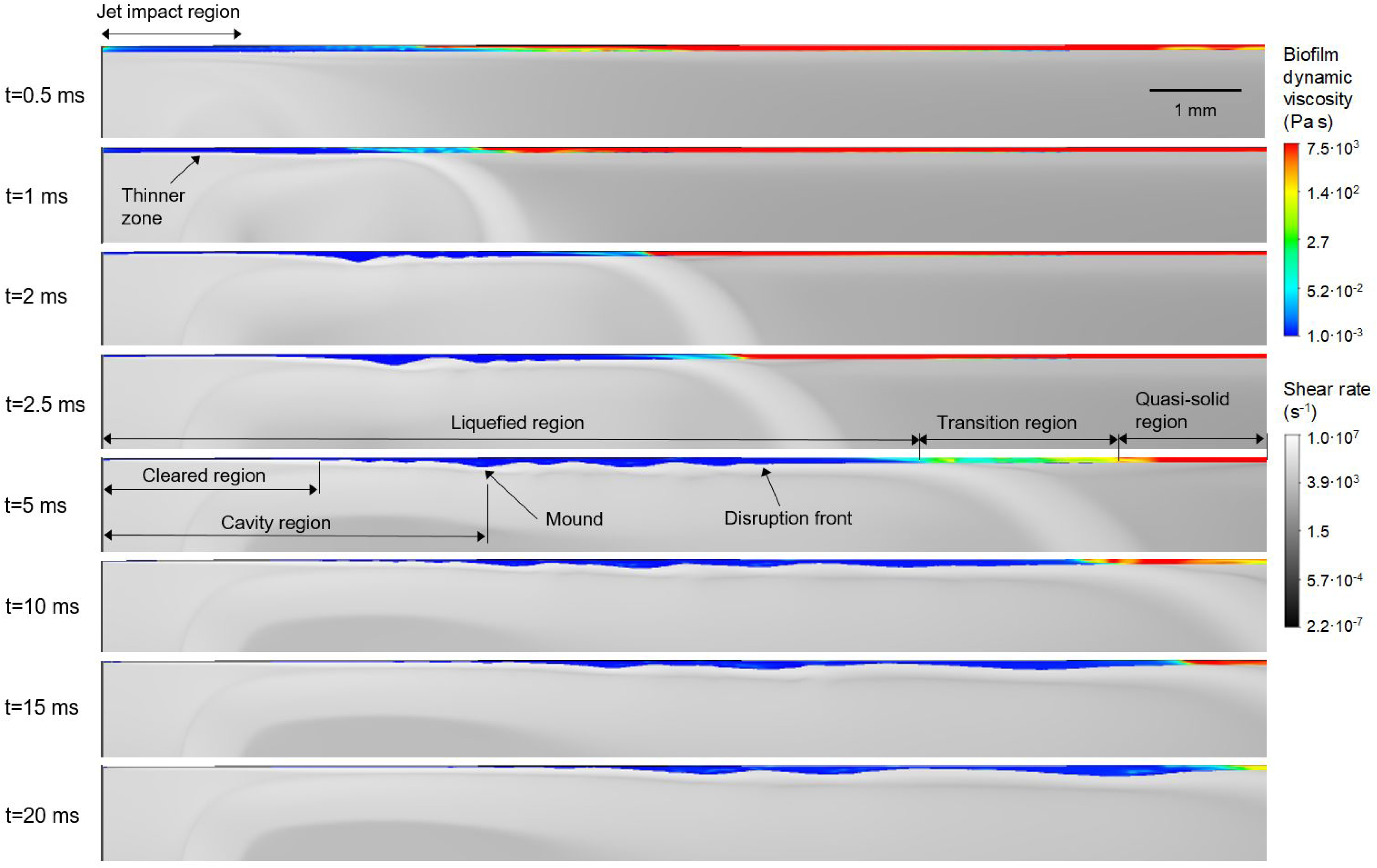
Simulated distributions of biofilm dynamic viscosity (color-scale area) and shear rate (gray-scale area) in the biofilm-disrupted region at different times (*η*=EVC_3_, *γ*=36 mN·m^-1^). Both biofilm viscosity and shear rate are displayed on logarithmic scales. (For interpretation of the references to color in this figure legend, the reader is referred to the Web version of this article.) An animation of the simulated biofilm rippling can be found in Supplementary Information **Video S5**.

The velocity component in the radial direction (*u_r_*, parallel with the initial biofilm surface) was dominant over the component in axial direction (*u_z_*), which indicated the drag direction on the biofilm surface (**Figure S4**). These shear forces and the pressure produced changes in biofilm thickness beginning from ∼0.7 ms in the impact zone. Slightly uneven biofilm surface can be observed in the area disrupted by the jet (Figure 7 and **Figure S3**), thus both pressure and shear stress forces initiate biofilm movement. Between 1 and 2 ms, the first biofilm ripples started to form. The ripple formation coincided with a larger biofilm area being liquefied, from a radius of 5 mm liquefied at 2 ms to about 8 mm at 5 ms. Interestingly, the movement of this part of fluidized biofilm produced pressure oscillations within the biofilm (**Figure S3**), thus generating the ripples. The largest gradients of pressure were observed for 5 ms, and decreasing further when the ripples were near a steady-state (from 15 ms). To characterize the biofilm ripples, the wavelength, characteristic frequency and ripples velocity were determined for both experimental and simulated data (*η*=EVC_3_, *γ*=72 and 36 mN·m^-1^). The values averaged in time are listed in Table 3, showing good agreement between simulated and experimental results.

**Table 3.** Ripples characterization: average wavelength (λ_R_), frequency (f) and average ripples velocity (v_R_) and their standard deviation.

| **Data** | **$\lambda_R$ (mm)** | **f (Hz)** | **$u_R$ (mm·s<sup>-1</sup>)** |
| --- | --- | --- | --- |
| Experimental | 0.9 ± 0.3 | 367 | 330 ± 110 |
| Simulated ($\eta$=EVC<sub>3</sub>, $\gamma$=72 mN/m) | 1.012 ± 0.13 | 311 | 315 ± 40 |
| Simulated ($\eta$=EVC<sub>3</sub>, $\gamma$=36 mN/m) | 1.047 ± 0.14 | 383 | 401 ± 54 |

### Sensitivity analysis

A sensitivity analysis was performed to determine the implications of the different model parameters in the biofilm disruption strategies, analyzing the biofilm displacement (Figure 8) and the development of the biofilm-cleared zone and the surface instabilities (Figure 9) both over jet exposure time. Parametric simulations were performed with changes in air velocity, biofilm thickness, biofilm viscosity and air-biofilm surface tension.

**Figure 8.**
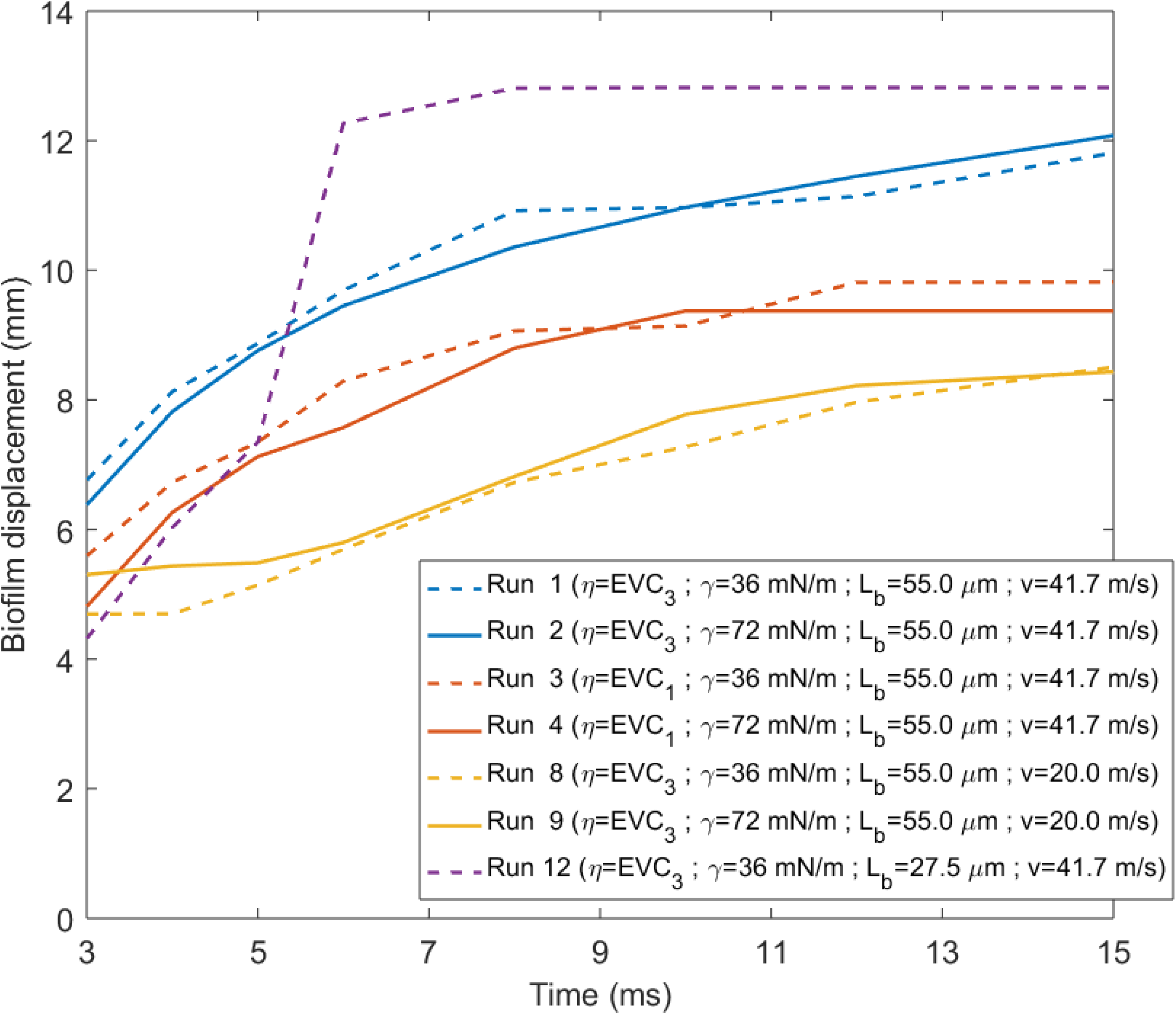
Parametric study of the biofilm displacement (mm) over jet exposure time (ms) for different model parameters: biofilm viscosity curves (run 1 and 2 vs run 3 and 4); jet velocities (run 1 and 2 vs run 8 and 9), biofilm thickness (run 1 vs run 12). Solid lines indicated the simulations computed with *γ*=72 mN·m^-1^, and dashed with *γ*=36 mN·m^-1^. (For interpretation of the references to color in this figure legend, the reader is referred to the Web version of this article.)

**Figure 9.**
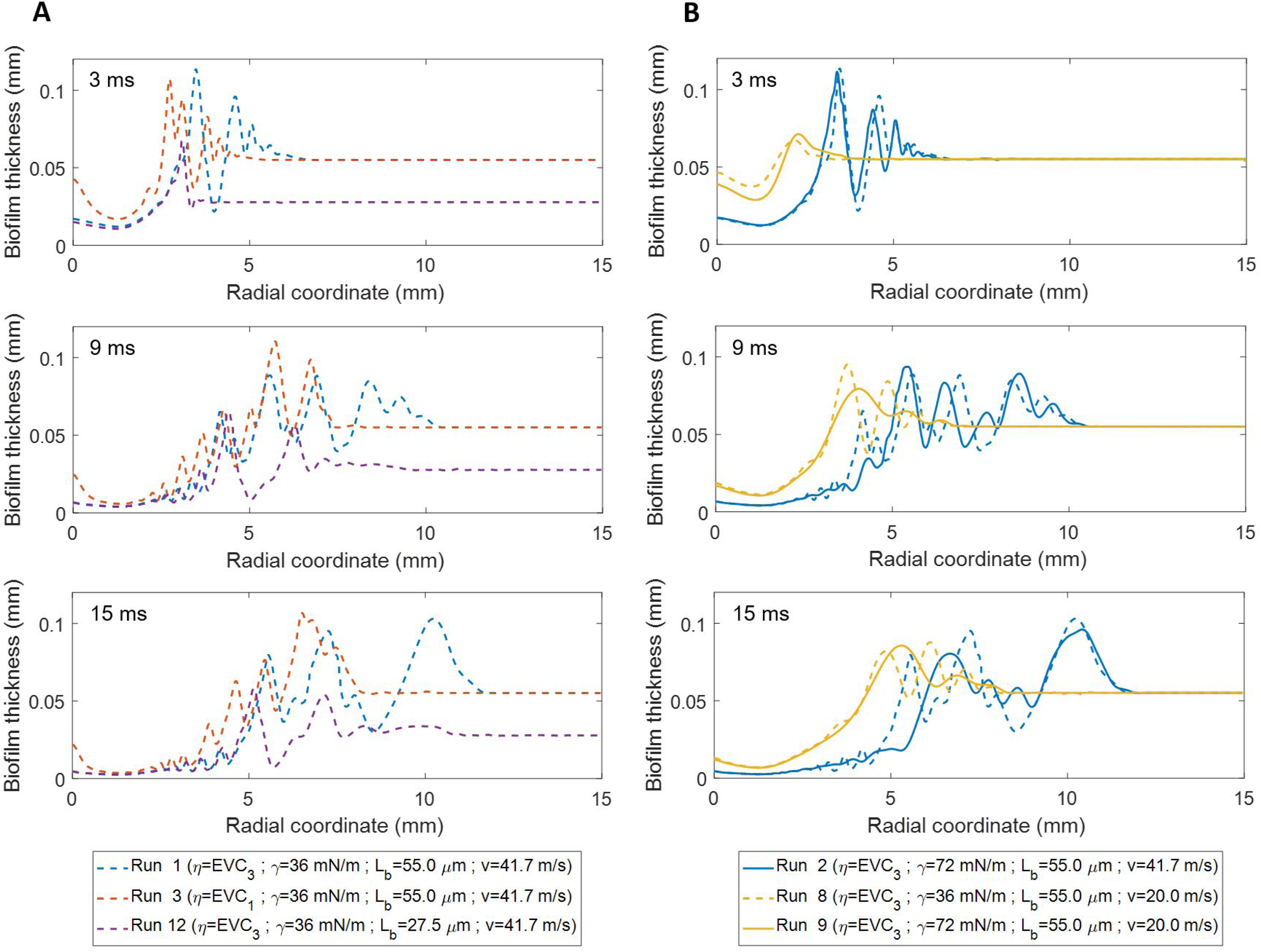
Sensitivity analysis data for the biofilm disruption produced by air-jet impingement for different model parameters: A) biofilm viscosity functions EVC (run 1 vs run 3), and biofilm thickness *L_b_* (run 1 vs run 12); B) jet velocities *v* (run 1 vs 8 and run 2 vs 9). (For interpretation of the references to color in this figure legend, the reader is referred to the Web version of this article.

#### Biofilm viscosity

Biofilms with higher viscosity (*η*=EVC_1_, runs 3 and 4) underwent smaller biofilm displacement due to the higher resistance to flow (Figure 8). Values below *η*=EVC_3_ (runs 1 and 2) meant a softer structure, disrupting the full biofilm length after only a few ms of jet exposure while experimentally the biofilm displacement was less than 6 mm for 2 ms. The biofilm with the lowest viscosity (*η*=EVC_3_, run 1) was disrupted over the largest radius, while biofilm residues remained unremoved in the cavity centre for the more viscous biofilm (*η*=EVC_1_, run 3) (Figure 9). Thus, expectedly, low values of biofilm viscosity lead to greater biofilm displacement and removal.

#### Surface tension

A lower surface tension (*γ*=36 mN·m^-1^, runs 1 and 3) allowed for quicker biofilm displacement initially (Figure 8) than with *γ*=72 mN·m^-1^ (runs 2 and 4), until arriving at similar steady-state displacement, possibly explained by the stabilization effect of surface tension. Moreover, the lower surface tension produced ripples with higher frequency and higher amplitude, i.e. the biofilm surface is more unstable (Figure 9). However, the cavity depth was not affected by the surface tension in the range of analyzed values.

#### Jet velocity

The largest and fastest displacements were achieved with high velocity (runs 1 and 3) due to the higher shear rates produced (Figure 8). For low velocity (runs 8 and 9), the biofilm started to move 5 ms later than for high velocity because the slower air jet reached the biofilm with the corresponding delay, generating also a smaller biofilm cavity (Figure 9B). Additionally, at *v*=60 m·s^-1^ the biofilm was removed much faster with full-length disruption after just 2 ms. In general, as the gas jet velocity increased the central biofilm cavity got deeper and wider, with the rim of the cavity rising above the original biofilm level, while the biofilm surface became more unstable, suffering larger surface perturbations.

#### Biofilm thickness

The thinner biofilm (*L_b_*=27.5 μm, run 12) was moved faster by the air jet and, consequently, reached stationary state sooner, in less than 8 ms, compared with >20 ms for the thicker film (*L_b_*=55 μm, run 1) (Figure 8). The thinner biofilm appeared slightly more stable than the thicker one, displaying less ripples (Figure 9A). Possibly, by having less material to be displaced favored the thin biofilm reaching quicker the steady-state. In addition, the cavity shape was very similar for the different biofilm thicknesses analyzed.

## DISCUSSION

### Disruption dynamics

Three distinct phases were identified during biofilm disruption by analysing the development of the biofilm ripple patterns. In the first phase from 1 to 6 ms (Figures 4 A,D), the ripples had a relatively regular wavelength and amplitude for the first wave formed, followed by a series of smaller waves until total wave decay at the disruption front. In the second phase from 7 to 14 ms (Figures 4 B,E) the waves appeared distorted, with a reduced amplitude and a more constant wavelength over the disruption area (i.e. the initial, smaller waves on the tail grow larger). Finally, from 15 to 20 ms (Figures 4 C,F) there were fewer ripples but with larger wavelengths than in the previous phases. Particularly, such wavelengths and ripples velocity characterizing perpendicular impingements were similar with those measured on *S. mutans* exposed to air-jets applied parallel to the surface (3).

The ripples dynamics produced by turbulent flow over biofilms has been previously related to the viscoelastic nature of biofilms (5, 7, 42), suggesting that the biofilm mechanical response is dominated by the EPS matrix properties (6, 43). Klapper et al. (6) hypothesized that the EPS matrix responds to stress by exhibiting: firstly, an elastic tension caused by the combination of polymer entanglement and weak hydrogen bonding forces; secondly, a viscous damping, where energy is absorbed as the biofilm flows and acts like a shock absorber due to the polymeric friction and hydrogen bond breakage; and thirdly, the polymers alignment in the shear direction, possibly leading to a shear-thinning effect in which the viscosity is lowered as the polymeric network structure of the biofilm matrix breaks down. The elastic tension may be related to the first disruption phase, where waves with similar pattern were generated in response to the initial jet-impingement. The viscous damping could correspond to the second disruption phase with distorted ripples. The polymers alignment could be associated with the last phase, where practically the ripples stop moving and changing form, being near to reaching steady-state. Possibly, the wave decay (where the ripples died down) occurred because the energy transmitted from the air to the biofilm at this distance and time was less than the viscous dissipation in the biofilm (44), being balanced by biofilm internal cohesive forces.

Interestingly, biofilm displacement reached a quasi-steady-state as the air jet velocity also approached the steady-state. This suggests that the relaxation time of air flow was on the same order as the plastic relaxation time of the biofilm. However, a relatively slow continuation of the biofilm movement after the air flow reached the quasi-stationary regime indicated the continuation of viscous damping within the biofilm.

Some model assumptions may compromise the accuracy of the determined ripples patterns. Assuming constant properties such as thickness and viscosity on the biofilm model has an effect on the ripples pattern formation. Further experimental techniques should be used to reveal heterogeneity in biofilm properties and to include them in future simulations. In addition, an accurate evaluation of the biofilm disruption should be done with 3D models. The difficulty however is not only the definition of 3D geometry and properties into the computational model, but more significantly a large increase in computing time.

### Conditions promoting disruption

Our model suggests that the air-jet exposures generating high shear rates, coupled with the biofilm shear-thinning behavior, produced rapid (within ms) biofilm disruption, thus affecting it both the biofilm properties and the applied forces intensity. The greatest interfacial instabilities are produced by the largest forces (i.e. high air-jet velocity) and the lowest values of biofilm properties (i.e. low viscosity, air-biofilm surface tension and thickness).

The similarity between the modeling and the experimental measurements suggests a non-Newtonian fluid behavior of *S. mutans* biofilms. Biofilm liquefaction, i.e. the complete breakdown of polymer inetwork nteractions in the biofilm matrix, is a mechanism that can explain the extremely quick disruptive effect induced by high shear air flows on the biofilms. The Herschel-Bulkley parameters of the estimated biofilm viscosity curve EVC_3_ described the required shear-thinning behavior, with the fitted yield stress (*σ_y_*=0.745 Pa) in accordance with the viscoelastic linearity limit (*σ*=3.5 Pa) previously determined by creep analysis (33). Biofilms in general show a mechanically viscoelastic behavior (42, 43), however the consistent results obtained considering the biofilm as a non-Newtonian fluid indicate that under turbulent flows the biofilms elastic behavior can be neglected, as recently reported (22). Additionally, biofilm expansion under non-contact brushing is attributed to its viscoelastic nature (45). Here, there was no evidence that the biofilm structure was expanded during impingement. Finally, although biofilm grown from a single species was analyzed here, the literature shows remarkable similarity in the viscoelastic response of many different types of biofilms when subject to shear stresses, even though the magnitude of the elastic and viscous moduli vary over many orders of magnitude(46), thus it is reasonable to conclude that biofilms formed from other species might exhibit similar flow behavior to described here, as suggested by the ripple patterns experimentally observed in *Pseudomonas aeruginosa* and *Staphylococcus epidermidis* biofilms exposed to high velocity shear flows (3).

Furthermore, the observed results highlight the importance of considering the correct representation of forces, which mechanically can disrupt biofilms. The numerical simulations indicated that inertial and interfacial tension forces are governing biofilm disruption by impact of turbulent air-jets, as the fluid dynamic activity reported for microdroplets sprays and power toothbrushes (27, 45). Specifically, the cavity formation, i.e. size and geometry, depends on a force balance at the free surface (23), thus assuming in our case that inertial forces controlled the cavity depth, while the cavity width was determined by both inertia and surface tension, indicated by the presence of small-amplitude ripples at the cavity edge. Ripples are produced because of pressure and shear stress variations in the gas in phase with the generated wave (44). For very thin fluid films, the fluctuations in the fluid have much larger components in the tangential direction than in the normal direction, consequently, the shear stress is the dominant mechanism (47), as showed in our case, in which was also identified a significant role for pressure variations. Therefore, two mechanisms were determined to produce moving biofilm ripples as a result of air jet impingement: 1) pressure oscillations generate biofilm ripples and 2) friction forces drag the biofilm along the support surface. Lastly, the simulation also revealed the important role of interfacial tension forces in the formation of surface instabilities, with less surface tension leading to more rippling (i.e. higher frequency and velocity). These results are in agreement with the possibly lower surface tension for biofilm-air (*γ*=36 mN·m^-1^) than for water-air (*γ*=72 mN·m^-1^), as measurements for *Bacillus subtilis*, *Pseudomonas fluorescens* and *P. aeruginosa* biofilms indicated values within the range 25-50 mN·m^-1^ (40, 48, 49). The amphiphilic character and surfactants production are associated as main effects controlling surface tension in microbial colonies and biofilms (49, 50), being attributed the surface tension reduction to the presence of surfactants (40, 43, 49). Biofilm surface tension differences could also explain the different ripple patterns observed between *S. mutans* biofilms and biofilms grown from *Pseudomonas aeruginosa* and *Staphylococcus epidermidis* (3). A lower surface tension intensified the formation of small-amplitude waves near the impact zone (i.e. the more “flexible” interface was more wrinkled). This increase in air-biofilm interfacial area could enhance the friction to flow. Moreover, there could be implications on the mass transfer: wavy interfaces will distort the diffusion boundary layer, and interfacial waves have been related to mass transfer enhancement (51, 52).

These results contribute to the existing knowledge of mechanisms promoting disruption though mechanical forces, opening new ways to optimize biofilm control strategies which rely on fluid shear. The model can be used to determine optimal parameters (e.g. jet velocity and position, angle of attack) to remove, or predict the spread of, the biofilm in specific applications (e.g. in dental hygiene or debridement of surgical site infections). The developed model also has potential application in predicting drag and pressure drop caused by biofilms in bioreactors, industrial pipelines and ship hull surfaces, as well as predicting how shear might influence the removal or spread of biofilms associated with medical devices such as orthopedic implants, voice prostheses, catheters and vascular stents.

## Acknowledgements

This work was financially funded in part by C15-69802-C2-2-R (MINECO/FEDER, UE) project. L. Prades was supported by grant BES-2013-066873 (FPI-2013, MINECO). P. Stoodley and S. Fabbri were supported in part by EPSRC DTP EP/K503130/1 award and in part by Philips Oral Healthcare, Bothell, WA, USA.

## SUPPLEMENTAL MATERIALS

### LIST OF TEXT FILES

**1) Computational and experimental investigation of biofilm disruption dynamics induced by high velocity gas jet impingement.**

### LIST OF FIGURES

**Figure S1.** Details of the defined mesh in the computational domain. A refined mesh was defined in the region ACGI satisfying the requirement y^+^≈1. A mesh growth rate no higher than ≈1.2 was used from the refined region to mesh the remaining domain. Mesh quality was checked with the orthogonal quality parameter, which had average values of 1 in all domain, confirming the good quality of the defined regular mesh.

**Figure S2.** Steady-state of the biofilm disruption after perpendicular air-jet impingement. The ripples die out when the biofilm flowed to the cleared space edge after ∼350 ms of jet exposure.

**Figure S3.** Computed pressure fields in the disrupted region (rectangle marked in Figure 6) for different times, i.e. 0.7, 0.8, 0.9, 1, 2, 2.5, 5, 10, 15, and 20 ms. Simulations were performed with *η*=EVC_3_ and *γ*=36 mN·m^-1^. Color scale: pressure in Pa. Larger pressures were in the air-jet impact zone. Pressure gradients formed in the in the biofilm phase were observed from t=2 ms.

**Figure S4.** Computed air-jet velocity profiles in X and Y directions (velocity u_r_ and u_z_, respectively) in the disrupted region (see Figure 6) for different times, i.e. 0.7, 0.8, 0.9, 1, 2, 2.5, 5, 10, 15, and 20 ms. Simulations were performed with *η*=EVC_3_ and *γ*=36 mN·m^-1^. Color scale: Velocity in m·s^-1^. Larger velocities in the gas phase were produced in the tangential direction to the biofilm phase, reaching maximum values of 45 m·s^-1^.

### LIST OF VIDEOS

**Video S1.** High-speed (2000 fps) recording of the air jet impingement experiment on a *S. mutans* biofilm attached to a glass surface, at an air velocity of 41.7 m·s^-1^, with a nozzle diameter of 2 mm. The biofilm consisted or larger clusters (lighter patches) distributed heterogeneously and separated by a base biofilm (grey). In the first few frames the shutter can be seen being lifted out of the way exposing the biofilm to the air jet. Biofilm ripples are forming radially away from the central impact area. After approximately 200 ms the biofilm had been “pushed” from the central area to from a cleared area (darker) of approximately 1.5 cm. This video was used to generate data for the computational analysis.

**Video S2.** Simulated biofilm ripples formation in time (X-Y top view), from a biofilm thickness of 55 μm with viscosity model EVC3 and surface tension γ=36 mN·m^-1^, subjected to a perpendicular air jet with velocity of 41.7 m·s^-1^.

**Video S3.** Simulated changes of biofilm thickness in time (0 – 20 ms) over the radial direction (X-Z lateral view) for viscosity model parameters EVC_3_ and surface tension *γ*=36 mN·m^-1^.

**Video S4.** Simulated biofilm ripples formation over time (X-Z lateral view), from a biofilm thickness of 55 μm with viscosity model EVC_3_ and surface tension *γ*=36 mN·m^-1^, subjected to a perpendicular air jet with velocity of 41.7 m·s^-1^. Colored surface: velocity magnitude (m·s^1^); Gray surface: biofilm area. In this simulation the jet is coming from the bottom of the screen and the biofilm is on the top.

**Video S5.** Simulated biofilm ripples formation over time (X-Z lateral view), from a biofilm thickness of 55 μm with viscosity model parameters EVC_3_ and surface tension *γ*=36 mN·m^-1^, subjected to a perpendicular air jet with velocity of 41.7 m·s^-1^. Gray-scale surface: shear rate (s^-1^); Color-scale: biofilm dynamic viscosity (Pa·s). In this simulation the jet is coming from the bottom of the screen and the biofilm is on the top.

